# The human microbiota is associated with cardiometabolic risk across the epidemiologic transition

**DOI:** 10.1101/595934

**Authors:** Na Fei, Beatriz Peñalver Bernabé, Louise Lie, Danny Baghdan, Kweku Bedu-Addo, Jacob Plange-Rhule, Terrence E. Forrester, Estelle V. Lambert, Pascal Bovet, Neil Gottel, Walter Riesen, Wolfgang Korte, Amy Luke, Stephanie A. Kliethermes, Brian T. Layden, Jack A. Gilbert, Lara R. Dugas

## Abstract

Oral and fecal microbial biomarkers have previously been associated with cardiometabolic (CM) risk, however, no comprehensive attempt has been made to explore this association in minority populations or across different geographic regions. We characterized gut- and oral-associated microbiota and CM risk in 655 participants of African-origin, aged 25-45, from Ghana, South Africa, Jamaica, and the United States (US). CM risk was classified using the CM risk cut-points for elevated waist circumference, elevated blood pressure and elevated fasted blood glucose, low high-density lipoprotein (HDL), and elevated triglycerides. Gut-associated bacterial alpha diversity negatively correlated with elevated blood pressure and elevated fasted blood glucose. Similarly, gut bacterial beta diversity was also significantly differentiated by waist circumference, blood pressure, triglyceridemia and HDL-cholesterolemia. Notably, differences in inter- and intra-personal gut microbial diversity were geographic-region specific. Participants meeting the cut-points for 3 out of the 5 CM risk factors were significantly more enriched with Lachnospiraceae, and were significantly depleted of Clostridiaceae, Peptostreptococcaceae, and *Prevotella*. The predicted relative proportions of the genes involved in the pathways for lipopolysaccharides (LPS) and butyrate synthesis were also significantly differentiated by the CM risk phenotype, whereby genes involved in the butyrate synthesis via lysine, glutarate and 4-aminobutyrate/succinate pathways and LPS synthesis pathway were enriched in participants with greater CM risk. Furthermore, inter-individual oral microbiota diversity was also significantly associated with the CM risk factors, and oral-associated *Streptococcus, Prevotella*, and *Veillonella* were enriched in participants with 3 out of the 5 CM risk factors. We demonstrate that in a diverse cohort of African-origin adults, CM risk is significantly associated with reduced microbial diversity, and the enrichment of specific bacterial taxa and predicted functional traits in both gut and oral environments. As well as providing new insights into the associations between the gut and oral microbiota and CM risk, this study also highlights the potential for novel therapeutic discoveries which target the oral and gut microbiota in CM risk.

## Introduction

Metabolic syndrome and cardiometabolic (CM) risk are associated with increased morbidity and mortality(1–3), and includes five risk factors: visceral obesity, elevated fasted blood glucose and elevated blood pressure, decreased high density lipoprotein (HDL) cholesterol, hypertriglyceridemia(4). In the US, as many as 35% of all US adults present with CM risk(5), when defined using the “Adult Treatment Panel criteria”(6). Between 1988 and 2012, the greatest increase in the prevalence of CM disease was seen among black men, estimated to be currently around 55%, and rose by 41% among black women(5). There is increasing evidence that the composition and metabolic function of the gut microbiota correlate with the progression and pathogenesis of CM disease, although the causality remains unclear(7).

One of the hallmarks of CM-related disease is the concomitant presence of low-grade systemic inflammation(8). Likewise, reduced intestinal bacterial diversity has also been reported in conditions with chronic inflammation and CM risk, with effects on the immune system(9, 10). The relative abundance of specific bacterial taxa has been shown to differ between individuals with CM risk to healthy participants. For instance, individuals with obesity, diabetes and cardiovascular disease, often have a significant reduction in fecal-associated short-chain fatty acid (SCFA)-synthesizing bacteria such as *Bifidobacteria, Roseburia, Faecalibacterium prausnitzii* and *Akkermansia muciniphilia*(11–13). All of these microorganisms have been shown to exhibit anti-inflammatory effects(13–16). Potential opportunistic pathogens, such as *Staphylococcus aureus* and those from the family Enterobacteriaceae, are often enriched in the stools of individuals with CM risk(17, 18). It has been suggested that these pro-inflammatory taxa contributes to the evolution of CM risk(17, 18).

There are several metabolic mechanisms through which the gut microbiota may contribute to the CM risk. Lipopolysaccharide (LPS) is a known precursor for the development of obesity and insulin resistance(19, 20) and an increase in the relative proportion of the bacteria that produce LPS in the gut has been associated with elevated systemic LPS concentration, likely resulting in high inflammation. Trimethylamine N-oxide (TMAO), which derived from microbial trimethylamine metabolism, is also increased in the stool of individuals with elevated CM risk(21, 22), although some find the inverse to be true(23) and one study found an inverse association with cardiac death in African Americans(24). Similarly, SCFAs such as propionate, acetate and butyrate produced by bacterial fermentation of indigestible fibers, are known to regulate host energy intake, expenditure and storage, decreases their concentrations have been associated with elevated CM risk(25, 26). Finally, specific gut bacteria have also been associated with altered bile acid composition, which also seem to play an important role in diabetes, obesity, non-alcoholic fatty liver disease and other metabolic diseases via the farnesoid X receptor (FXR) and G protein-coupled bile acid receptor (GPCR) signaling pathway(27–29). These metabolic pathways represent mechanisms through which the gut microbiota may influence CM risk, and by which they might also serve as potential therapeutic targets for treating elevated CM risk.

Emerging data suggests that the oral microbiota may also be linked to elevated CM risk(30–32). Oral bacteria, especially potential pro-inflammatory pathogens like *Pseudomonas* and *Enterobacter*, have been detected in human atherosclerotic plaques, associated with CM risk (31, 33). Hence, systemic inflammation triggered by oral pathogens may be an important component in the pathogenesis of systemic disease(30, 34). However, to date, it is unclear whether there is an association between the oral and gut microbiota and elevated CM risk, and whether the oral microbiota exhibits similar associations as gut microbiota does with CM risk. If this is the case, then, the oral microbiota, which is substantially more convenient to collect, may be a proxy to determine gut microbiota-derived associations that may have a more specific direct mechanistic association with elevated CM risk. However, a lack of large-scale cohort studies limits our interpretation of CM risk-microbiota associations, particularly across diverse human populations.

In this study, we leveraged African-origin participants enrolled in the “Modeling the Epidemiologic Transition Study” (METS)(35) cohort to determine the association between the gut (stool-derived) and oral (saliva-derived) microbiota and elevated CM risk. We characterized the gut- and oral-associated microbiota and indices of CM risk in 655 participants of African-origin, aged 25-45, from Ghana, South Africa, Jamaica, and the United States (US). The central hypothesis is that the oral microbial composition is associated with the gut microbial composition, and that both are associated with elevated CM risk in blacks.

## Results

### Participant characteristics

Previously 2,506 adults from Ghana, South Africa, Jamaica, The Seychelles, and US were recruited in 2009, and prospectively followed on an annual basis in METS(35). The current study included a subsample of 655 participants, of which 196 were Ghanaian, 176 were South African, 92 were Jamaican, and 191 were from the US (**Table 1**). Approximately 60% of the participants were women. The average age among all the participants was 34.9 ± 6.4 years, and participants from South Africa and Jamaica were significantly younger than US participants (p <0.001 and p=0.016, respectively). Men and women from Ghana, South Africa and Jamaica weighed significantly less (p <0.001) than their US counterparts (63.2 ± 12.0, 76.2 ± 20.5, and 80.0 ± 21.1 kg, respectively), and there were significantly more US men and women that were overweight and obese (81.3%, p <0.001 for all) compared with the other cohorts (i.e., Ghana, 33.2%; South Africa, 55.7%; Jamaica 65.2%) (**Table 1**). Americans also slept the least number of nightly hours (6.7 ± 1.4 hrs) compared to Ghanaians (7.9 ± 1.4 hrs, p<0.001), Jamaicans (7.3 ± 2.1 hrs, p<0.033), and South Africans (10.5 ± 1.7 hrs, p<0.001), and reported significantly higher prevalences of cigarette smoking (37.7%, p<0.001), and alcohol consumption (84.3%, p<0.001) compared to the other 3 sites, after adjusting for age, sex and BMI (**Table 1**).

**Table 1.** Participant characteristics by site. The US is the reference site.

|  | Ghana | South Africa | Jamaica | US | Overall |
| --- | --- | --- | --- | --- | --- |
| **N** | N=196 | N=176 | N=92 | N=191 | N=655 |
| **Age (y)** | 35.8 ± 6.6 | 33.3 ± 5.9* | 33.9 ± 6.2* | 36.0 ± 6.3 | 34.9 ± 6.4 |
| **Weight (kg)** | 63.2 ± 12.0** | 76.2 ± 20.5** | 80.0 ± 21.1** | 94.2 ± 25.0 | 78.1 ± 23.3 |
| **Height (cm)** | 161.7 ± 7.6** | 163.7 ± 7.7** | 166.8 ± 10.9** | 169.6 ± 8.4 | 165.2 ± 9.0 |
| **BMI (kg/m<sup>2</sup>)** | 24.3 ± 5.0** | 28.7 ± 8.2** | 29.1 ± 8.9** | 32.8 ± 8.8 | 28.6 ± 8.4 |
| **Normal weight (BMI<25 kg/m<sup>2</sup>), N, %** | 131, 66.8%** | 78, 44.3%** | 32, 34.8%** | 36, 18.9% | 277, 42.3% |
| **Overweight (BMI≥25 kg/m<sup>2</sup>-<30 kg/m<sup>2</sup>)** | 41, 20.9% | 28, 15.9%** | 28, 30.4% | 52, 27.2% | 149, 22.8% |
| **Obese (BMI≥30 kg/m<sup>2</sup>)** | 24, 12.2%** | 70, 39.8%** | 32, 34.8%** | 103, 53.9% | 229, 35.0% |
| **Fat-free mass (kg)** | 44.3 ± 7.4** | 44.6 ± 7.2** | - | 55.1 ± 11.2 | 48.1 ± 10.2 |
| **Fat mass (kg)** | 19.0 ± 9.2** | 31.7 ± 16.2** | - | 39.3 ± 18.4 | 29.9 ± 17.3 |
| **% Body fat** | 29.2 ± 9.84** | 39.3 ± 11.0 | - | 39.9 ± 10.8 | 36.0 ± 11.6 |
| **Sleep hours (hrs/night) \*** | 7.9 ± 1.4** | 10.5 ± 1.7** | 7.3 ± 2.1* | 6.7 ± 1.4 | 8.2 ± 2.2 |
| **Smokers\*** | 4, 2.0%** | 48, 27.3 %** | 10, 10.9%** | 72, 37.7% | 134, 20.5% |
| **Drinkers\*** | 59, 30.6%** | 75, 42.9%** | - | 161, 84.3% | 295, 52.8% |

\*p<0.05, \*\*p<0.01 compared to US.
\*Adjusted for age, sex and BMI

### Cardiometabolic risk

We used the National Cholesterol Education Program’s Adult Treatment Panel III (NCEP/ ATP III) criteria for metabolic syndrome(6) to indicate CM risk, and as follows; waist circumference >102 cm in men and >88 cm in females; elevated blood pressure (≥130/85 mm Hg), or receiving treatment for hypertension; hypertriglyceridemia (≥150 mg/dL) or receiving treatment; low high-density lipoprotein (HDL) cholesterol (<40 mg/dL in males and <50 mg/dL in females), or receiving treatment; and elevated fasting plasma glucose (≥110 mg/dL) or receiving treatment for type 2 diabetes. Individuals with a BMI ≥25 kg/m^2^ were classified as overweight, and ≥30 kg/m^2^ were categorized as obese(36). For the purpose of our analysis, we dichotomized CM risk creating a CM risk phenotype, whereby those with elevated CM risk had at least 3 of the five CM risk factors, compared to those with 2 or less. Because the Jamaican participants were missing HDL and triglyceride concentration measures in blood, they were excluded from the overall CM risk analysis. This resulted in a total of 563 participants with complete CM risk data. Overall, 46 of these 563 participants (8.2%) presented with elevated CM risk. Compared to the US participants (14.7%), the prevalence of elevated CM risk was significantly lower among the Ghanaians (1.5%, p<0.001), and trended lower among the South Africans (8.5%, p=0.09) (**Table 2**).

**Table 2.** Cardiometabolic risk by site, and overall. The US is the reference site, and comparisons are adjusted for age, sex and BMI.

|  | **Ghana** | **South**<br>**Africa** | **Jamaica** | **United**<br>**States** | **Overall** |
| --- | --- | --- | --- | --- | --- |
|  | N=196 | N=176 | N=92 | N=191 | N=655 |
| **Fasted plasma glucose**<br>**(mg/dL)** | 100.7 ±<br>12.2 | 83.3 ±<br>18.1** | 99.6 ±<br>15.5 | 106.2 ±<br>40.4 | 97.5 ±<br>26.8 |
| **Elevated fasted plasma**<br>**glucose, N, %** | 2, 1.0%** | 8, 4.6%** | 18, 19.6% | 45, 23.9% | 73, 11.2% |
| **Systolic blood pressure**<br>**(mmHg)** | 112.9 ±<br>13.5** | 124.4 ±<br>20.5* | 112.7 ±<br>12.5** | 123.3 ±<br>17.6 | 119.0 ±<br>17.5 |
| **Diastolic blood pressure**<br>**(mmHg)** | 66.6 ±<br>11.0** | 79.4 ±<br>12.8 | 72.0 ±<br>34.0** | 80.5 ± 13.1 | 74.9 ±<br>18.1 |
| **Elevated blood pressure,**<br>**N, %** | 6, 3.1%** | 26, 14.8% | 1, 1.1%* | 25, 13.1% | 58, 8.9 |
| **Triglycerides (mg/dL)** | 82.1 ± 36.5 | 88.2 ±<br>55.1 | - | 98.5 ± 62.1 | 89.5 ±<br>52.5 |
| **Hypertriglyceridemia, N, %**<br>**(N=563)** | 12, 6.2% | 16, 9.1% | - | 29, 15.7% | 57, 10.3% |
| **HDL-cholesterol (mg/dL)** | 46.1 ±<br>13.6** | 49.9 ±<br>15.0* | - | 52.1 ± 15.1 | 49.3 ±<br>14.7 |
| **Low HDL-cholesterol, N, %**<br>**N=563** | 104,<br>53.6%** | 79, 45.4%* | - | 64, 34.8% | 247,<br>44.8% |
| **LDL-cholesterol (mg/dL)** | 100.7 ±<br>29.1 | 92.9 ±<br>32.6** | - | 110.1 ±<br>35.5 | 101.4 ±<br>33.1 |
| **Total cholesterol (mg/dL)** | 163.3 ±<br>34.3** | 160.1 ±<br>36.1** | - | 183.0 ±<br>38.6 | 168.9 ±<br>37.7 |
| **High waist circumference**<br>**N, % N=563** | 37,<br>18.9%** | 78,<br>44.3%** | 37,<br>40.2%** | 116, 60.7 | 268,<br>40.9% |
| **Cardiometabolic disease**<br>**N, % N=563** | 3, 1.5%* | 15, 8.5% | - | 28, 14.7 | 46, 8.2% |

Cardiometabolic risk is defined by three out of five risk factors: waist circumference >102 cm in males and >88 cm in females; elevated blood pressure (≥ 130/85 mm Hg), or receiving treatment; hypertriglyceridemia (≥ 150 mg/dL), or receiving treatment; low HDL (<40 mg/dL in males and <50 mg/dL in females), or receiving treatment; and elevated fasting plasma glucose (≥ 110 mg/dL) or receiving treatment. \*p<0.05, \*\*p<0.01 compared to US.

### Gut-derived bacterial diversity associates with country of origin and CM risk factors

The associations between the gut-derived microbial diversity measured by 16S rRNA amplicon sequences using exact sequence variants (ESVs) and indices of elevated CM risk were calculated across the four cohorts. All analyses were adjusted for age, sex and BMI. Gut bacterial alpha diversity (intrapersonal gut diversity, measured by Shannon and Chao1 indexes)(37) was significantly greater among the Ghanaians and South Africans, compared to the US participants (p <0.001; **Figure 1**, **Supplementary Figure 1**). The Shannon index, which incorporates both the richness and evenness of the community, was lower among participants with elevated blood pressure from South Africa (p <0.05) and Ghana (p <0.01), but showed no significant relationship among the US participants or the Jamaicans (**Figure 1**, **Supplementary Figure 2**). Elevated fasted blood glucose was significantly associated with lower alpha diversity, but only among the Jamaicans (p <0.05; **Figure 1**, **Supplementary Figure 2**). Individuals with elevated waist circumference, triglyceride concentration and HDL levels showed no significant associations for gut microbial alpha diversity (adjusted for age, sex and BMI, p >0.05, **Supplementary Figure 2**). These results suggest that decreased gut microbial alpha diversity is associated some of the CM risk factors (i.e., elevated blood pressure and elevated fasting blood glucose), however, these associations are country specific.

**Figure 1.**
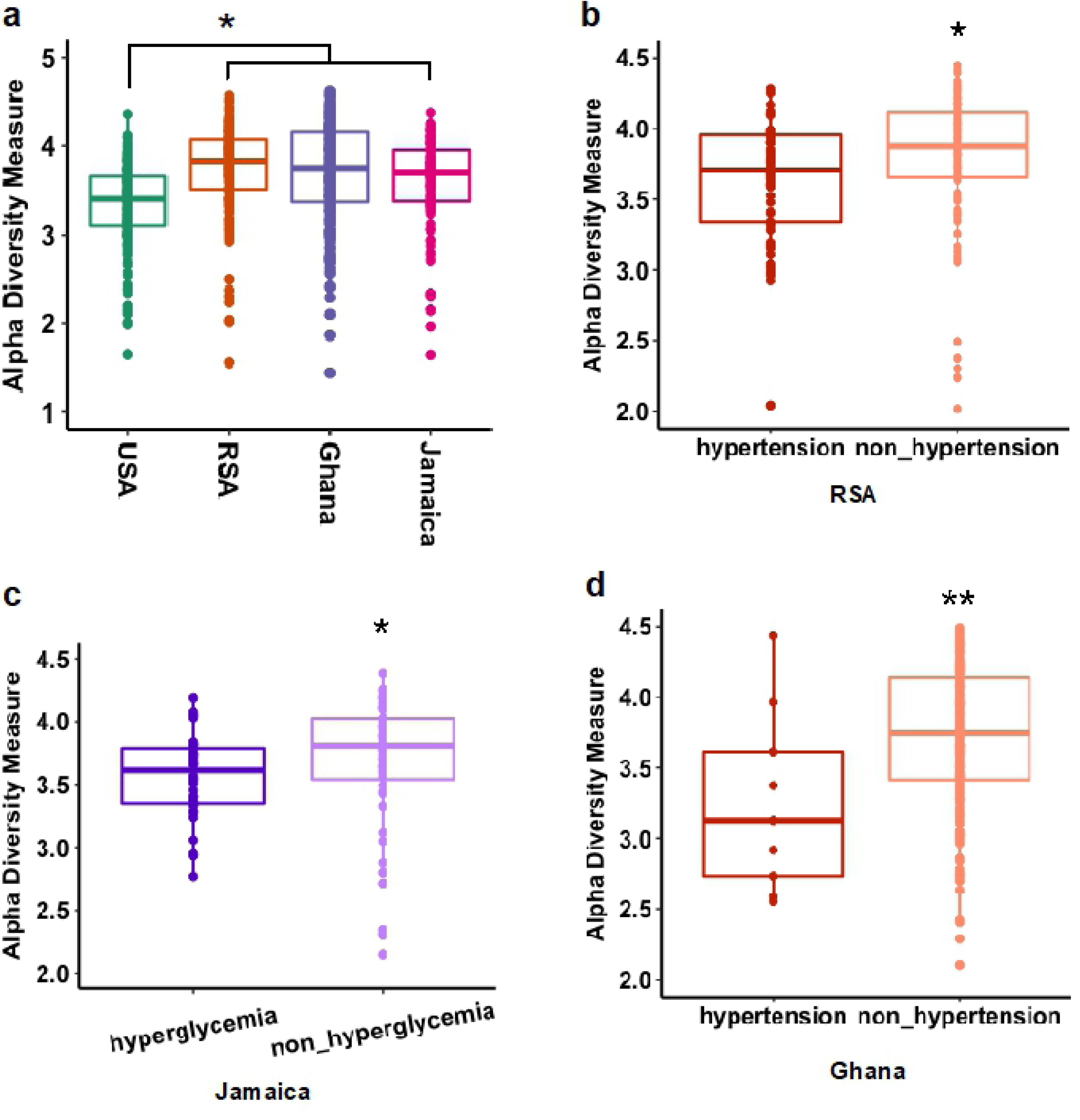
Gut bacterial alpha diversity associations with cardiometabolic risk factors. (adjusted for age, sex and BMI, only significant associated were shown here). **a**) Alpha diversity (Shannon index) in the four study sites (USA, RSA, Ghana and Jamaica); **b**) association with hypertension in RSA; **c**); association with elevated fasting blood glucose in Jamaica; **d)**; association with hypertension in Ghana. * p <0.05; ** p <0.01.

**Figure 2.**
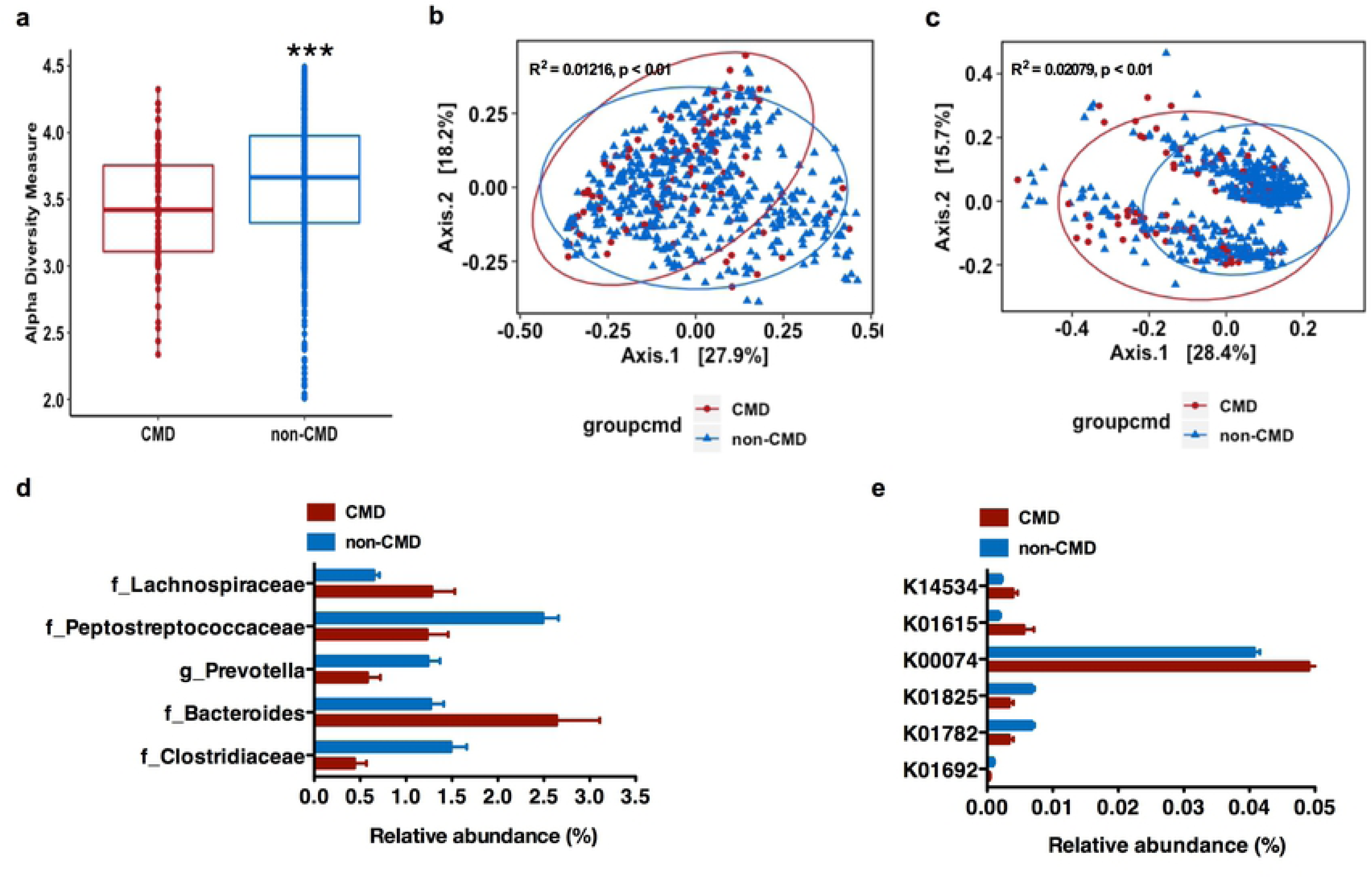
Gut bacterial structure and function correlates with cardiometabolic (CM) risk phenotype. (**a**), Alpha diversity analysis (Shannon Index) from 16S rRNA gene sequence data of gut microbiota against CM risk; (**b-c**), Principal coordinate analyses (PCoA) of weighted (**a**) and unweighted (**b**) UniFrac distance of gut microbiota composition against CM risk (fdr-corrected p <0.01); (**d**), Specific gut bacterial taxa are differentially abundant (in relative terms) between study sites with and without elevated CM risk (only significantly differential exact sequence variants (ESVs) with relative abundance ≥ 1% in at least one group shown. Data shown are means ± S.E.M.; p(fdr-corrected) <0.05); (**e**), Relative abundance of the genes involved in the four different pathways for butyrate synthesis against CM risk of gut microbiota across all study sites. (Data shown are means of percentages ± S.E.M. p(fdr-corrected) < 0.05). CMD means CM risk defined as at least 3 CM risk factors from five: waist circumference, elevated blood pressure, elevated blood fasting glucose, elevated triglyceride and low HDL concentration in USA, RSA, and Ghana *** p < 0.001. false discovery rate

Gut bacterial beta diversity was significantly different between countries [false discovery rate (fdr)-corrected p <0.01; **Supplementary Figure 3**]. Therefore, as with alpha diversity, we derived associations of beta diversity against CM risk within each country independently (**Supplementary Figure 4 and 5**). Weighted and unweighted UniFrac distances, which compute differences between microbial communities based on phylogenetic information(38), were significantly different between high and low waist circumference, but only among the South Africans (fdr-corrected p<0.05) and Ghanaians (fdr-corrected p<0.01). The same was true for elevated blood pressure, whereby weighted UniFrac was significantly different among the South Africans (fdr-corrected p <0.05) and Ghanaians (fdr-corrected p <0.05), but interestingly, unweighted UniFrac was only significantly different in the South Africa cohort (fdr-corrected p <0.05). This might suggest that the differences seen in the South Africans may be in part due to differences in the abundance of rare bacterial taxa, while in Ghana the differences may be due to more abundant bacterial taxa in the fecal samples. Similarly, for triglyceride concentrations, only weighted UniFrac was significantly different and only among the South Africans (fdr-corrected p <0.05), while for HDL concentrations only weighted UniFrac was significantly different among US participants (fdr-corrected p <0.05). Hyperglycemia was not significantly correlated with beta diversity in any of the cohorts. These results suggest that gut microbial structure is significantly associated with individual CM risk factors, and that these associations are geographically dependent, given that the contribution of the abundant versus rare bacterial taxa varied with country of origin. Taken together, contributions of CM risk as well as environmental factors, including country of origin, result in inter-individual dissimilarities in the gut microbial composition (**Supplementary Table 1**).

**Figure 3.**
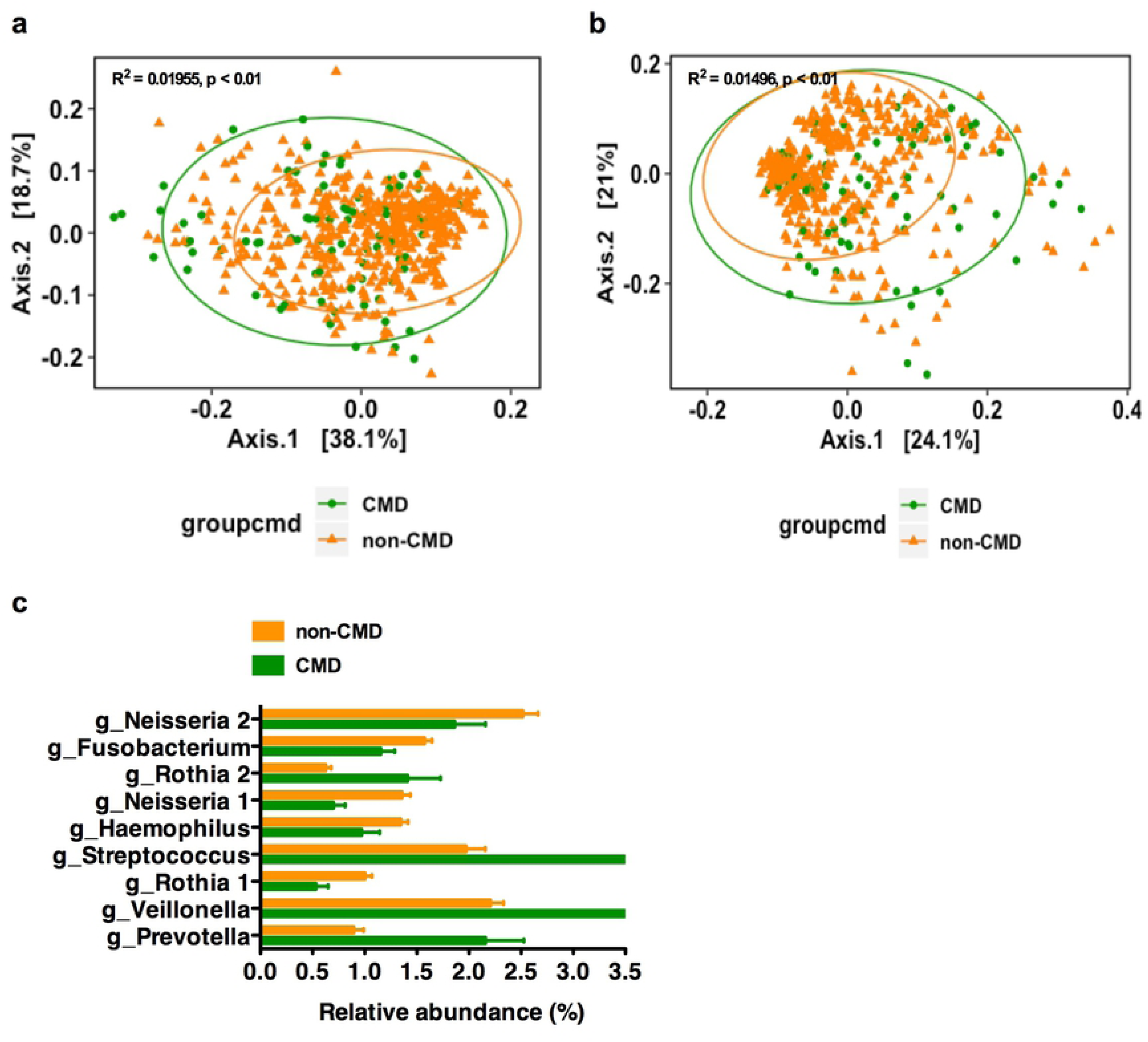
Oral-associated bacterial structure correlates with CM risk phenotype. (**a-b**), Principal coordinate analyses (PCoA) of weighted (**a**) and unweighted (**b**) UniFrac distance of oral microbiota composition against CM risk (fdr-corrected p <0.01). (**c**), Specific oral bacterial taxa are differentially abundant (in relative terms) between study sites with and without CM risk (only significantly differential exact sequence variants (ESVs) with relative abundance ≥ 1% in at least one group shown. Data shown are mean± S.E.M. p(fdr-corrected) < 0.05). CMD means CM risk, including at least 3 CM risk factors of waist circumference, elevated blood pressure, elevated blood fasting glucose, high triglyceridemia and low HDL-cholesterolemia in USA, RSA, and Ghana. fdr, false discovery rate.

### Gut microbial structure and function correlate with CM risk phenotype

We next investigated whether the gut microbiota correlated with an elevated CM risk phenotype in the participants from USA, Ghana and South Africa. Jamaican’s were excluded from this analysis due to missing lipid data, and analyses were adjusted for age, sex and BMI. Participants with elevated CM risk phenotype (at least 3 risk factors) had significantly lower fecal alpha diversity (measured by Shannon index), compared to those without elevated CM risk (p <0.001; **Figure 2a**). Furthermore, weighted and unweighted UniFrac distances were significantly different between the two CM risk phenotypes (fdr-corrected p < 0.05; **Figure 2b**, **2c**). We were able to identify the fecal bacterial ESVs which had significantly different abundance as a function of elevated CM risk. For these ESVs with a relative abundance ≥1% in at least one group (adjusted for country, sex, age and BMI), participants with elevated CM risk were significantly enriched with 2 bacterial ESVs annotated to family Lachnospiraceae and genus *Bacteroides*, while participants without elevated CM risk had were enriched in 3 bacterial ESVs annotated to family Clostridiaceae, Peptostreptococcaceae, and genus *Prevotella* (fdr-corrected p <0.05; **Figure 2d**).

Specific gut bacterial taxa were also associated with individual CM risk across the entire cohort (adjusted for country, age, sex and BMI, **supplementary Figure 6 and Supplementary Table 2**) or when stratified by sex (adjusted for country, age and BMI, **supplementary Figure 6 and Supplementary Table 2**). We found that the gut microbial taxa level was profoundly altered in participants for each of the CM risk factors. For example, among participants with an elevated waist circumference, hyperglyceridemia, elevated blood pressure or low HDL concentration, there were a significant enrichment of bacterial ESVs annotated to genera belonging to family Lachnospiraceae, Enterobacteriaceae and Clostridiaceae; genera *Streptococcus, Coprococcus* and *Blautia* (fdr-corrected p <0.05). Conversely, bacterial ESVs annotated to genus *Prevotella* (family Prevotellaceae), and family Enterobacteriaceae, Clostridiaceae, and Peptostreptococcaceae were also significantly enriched among participants without CM risk (fdr-corrected p <0.05; supplementary results). As expected, there were also several gut bacterial taxa that were differentially abundant between each country and each CM risk factor (**supplementary Figure 7 and Supplementary Table 2**).

The 16S rRNA amplicon data was used to predict gene abundance using Piphillin(39) to establish differences in the gut microbial metabolic potential in several well-known pathways associated with elevated CM risk, such as LPS biosynthesis, SCFA metabolism, TMA N-oxide (TMAO) biosynthesis, and genes associated with secondary bile acid biosynthesis. No statistically significant differences in the proportion of genes encoding components of TMA N-oxide (TMAO) biosynthesis, LPS biosynthesis, and secondary bile acid biosynthesis were observed among participants who had an elevated CM risk phenotype (at least 3 risk factors), compared to those without CM risk (adjusted for country, sex, BMI and age, p >0.05). However, the relative abundance of the genes involved in the four different pathways for butyrate synthesis showed that several genes that encode for enoyl-CoA hydratase enzymes (K01692, K01782, K01825) in the acetyl-CoA pathway were enriched in participants without elevated CM risk; while, crotonase (K01715) and β-hydroxybutyryl-CoA dehydrogenase (K00074) in the acetyl-CoA pathway were enriched in participants with elevated CM risk. Notably, the abundance of predictive genes that encode for glutaconyl-CoA decarboxylase (α, β subunits) (K01615) in the glutarate pathway and the 4-hydroxybutyryl-CoA dehydratase (K14534) in the 4-aminobytyrate/succinate pathway were enriched in participants with elevated CM risk (adjusted for country, sex, age and BMI, fdr-corrected p < 0.05; **Figure 2d**).

We also explored whether gut bacterial functions were associated with each of the 5 individual CM risk factor within each of the countries. Gut microbial predicted genes involved in LPS and SCFA biosynthesis pathways showed that genes involved in the butyrate synthesis via lysine, glutarate and 4-aminobutyrate/succinate pathways and LPS synthesis pathway were differentially enriched among the cohort with an elevated individual CM risk factor (**supplementary Figure 8-11**). Specifically, predicted genes involved in LPS biosynthesis pathways were enriched among participants with an elevated waist circumference in Ghana and US; participants with an elevated blood pressure in Ghana and Jamaica; participants with an elevated fasted blood glucose in Jamaica and the US, and those with low HDL in South Africa and the US. As the butyrate synthesis via lysine, glutarate and 4-aminobutyrate/succinate pathways, predicted genes involved in the glutarate pathway were significantly enriched in participants with a high waist circumference in South Africa and Ghana, and also participants with low HDL concentrations in the US and South Africa; predicted genes involved in the lysine pathway were enriched in participants with elevated blood pressure in Ghana; and predicted genes involved in the 4-aminobytyrate/ succinate pathway in South Africans were significantly associated with low HDL concentrations (see detail in supplementary results; **supplementary Figure 8-11**).

Generalized linear models (GLM) were applied to explore the associations between alpha diversity, bacterial taxa and predicted functional genes and total CM risk (captured as the sum of CM risk factors in each participant). For instance, there was a significant positive correlation between the proportion of KEGG ID K01615 in the glutarate pathway and K14534 in the 4-aminobytyrate/succinate pathway and total CM risk score (i.e. a z-score transformation to generate continuous discrete measurements, controlling for country, age, sex, and BMI, p <0.05, see methods for more details). However, there were no significant associations between diversity or differential bacterial taxa and total CM risk (p >0.05). To explore whether the gut microbiota predicted elevated CM risk (three out of the five risk factors), random forest regression was used to determine the gut bacterial ESVs against the total CM risk (z-score transformed) pooling the 3 sites with CM risk data. Elevated CM risk could be predicted by gut-associated ESVs and explained almost 15% of the elevated CM risk variance among those with CM risk data.

### Oral-associated bacterial diversity associates with country of origin and CM risk factors

On the other hand, oral microbiota diversity was associated with the country of origin, but not with the individual CM risk factors. Shannon diversity was significantly greater among the Ghanaians compared to the US participants (adjusted for age, sex and BMI, p <0.05; **Supplementary Figure 12**). As oral microbial alpha diversity was significantly different between countries, we correlated CM risk against alpha diversity for each country independently, and found that none of the CM risk factors were significantly associated with oral microbial alpha diversity (p >0.05, **Supplementary Figure 13**). However, oral microbial beta diversity was significantly different between participants from different countries (fdr-corrected p <0.01, **Supplementary Figure 14**), and therefore, tests of association were performed separately for each country. In the US sample, weighted UniFrac distance was significantly different between individuals with either high and normal waist circumference (fdr-corrected p <0.01) and similarly, between participants with either elevated and normal glucose concentrations (fdr-corrected p <0.05, **Supplementary Figure 15**). Unweighted UniFrac distances were significantly different between individuals with either elevated and normal waist circumference in the US (fdr-corrected p ≤ 0.01), Ghana (fdr-corrected p <0.05), and South Africa (fdr-corrected p <0.05) (**Supplementary Figure 16**). In the US alone, unweighted UniFrac was significantly different by HDL risk (fdr-corrected p <0.05) (**Supplementary Figure 16**). Therefore, oral beta diversity, and hence microbial structure, was significantly associated with several CM risk factors, but similarly differed by country, suggesting that environmental factors are critical for inter-individual dissimilarities in oral microbial composition (**Supplementary Table 3**).

### Oral-associated bacterial community structure correlates with the elevated CM risk phenotype

We next investigated whether the oral microbiota correlated with the elevated CM risk phenotype (3 out of five risk factors) and CM risk score in the US, Ghanaian and South African participants. Alpha diversity was not significantly different between low and elevated CM risk (adjusted for age, sex and BMI, p(fdr-corrected) <0.05, **Supplementary Figure 17**). However, both weighted and unweighted UniFrac distance were significantly differentiated by CM risk (fdr-corrected p <0.01, **Figure 3a**, **3b**). The differential taxa analysis (with relative abundance ≥1% in at least one group) showed that participants with elevated CM risk had a significant enrichment of 4 oral-bacterial ESVs annotated to the genera *Prevotella, Veillonella*, and *Streptococcus*; while participants without elevated CM risk had a significant enrichment of 5 bacterial ESVs annotated to genus *Haemophilus, Neisseria, Fusobacterium* (adjusted for country, sex, BMI and age, p(fdr-corrected) <0.05; **Figure 3c**). Several oral-bacterial ESVs annotated to the genus *Rothia* presented opposite behaviors (i.e., some were enriched, and some were depleted in participants with elevated CM risk compared with low CM risk)

At a more granular level, specific oral bacterial taxa were also associated with each CM risk factor across the entire cohort (adjusted for country, age, sex and BMI; **supplementary Figure 18 and supplementary Table 4**) or when stratified by sex (adjusted for country, BMI and age; **supplementary Figure 18 and supplementary Table 4**), separately. The results indicate that the oral microbiota of participants who have greater CM risk, e.g. a higher waist circumference, elevated blood pressure or low HDL concentration, are significantly enriched for potentially pro-inflammatory taxa, including *Streptococcus, Prevotella*, and *Veillonella* (fdr-corrected p <0.05; supplementary results). There were also specific oral bacteria that were differentially abundant between each country by each CM risk factor (fdr-corrected p <0.05; supplementary results; **supplementary Figure 19 and Supplementary Table 2**).

To determine whether the taxa could identify participants with elevated CM risk, we once again used GLM to determine the association between the differential proportional oral bacterial taxa and total CM risk (sum of the number of CM risk factors for each participant). There was a significantly positive association between the proportion of a *Streptococcus* ESV and total CM risk (z-score transformation), controlling for country, age, and sex p <0.05). To test whether the oral microbiota can identify participants with the elevated CM risk phenotype, random forest regression was used to examine the association of oral bacterial ESVs against CM risk (z-score transformed). Among participants with elevated CM risk, oral bacterial ESVs, accounted for almost 8% of the variance.

### Correlation of beta diversity between gut and oral microbiota

Finally, to explore whether the beta diversity trends seen in the oral microbiota correlated with those in the gut microbiota, we applied a Mantel test using both unweighted and weighted UniFrac distance matrices. Pooling all countries, there was a significant correlation with both unweighted (r = 0.094, p =0.001) and weighted (r =0.082, p =0.001) UniFrac distance between the gut and oral microbiota (**Supplementary Figure 20 and 21**). However, within each country, only the weighted UniFrac metric had a significant correlation between oral and gut microbial beta diversity, and only among the US and South African participants (p = 0.034, r = 0.073 and p = 0.021, r = 0.09 respectively) (**Supplementary Figure 20 and 21**).

## Discussion

In our study of African-origin adults from Ghana, South Africa, Jamaica and the US, we performed a comprehensive analysis exploring the relationship between the gut and oral microbiota and elevated CM risk, representing 4 geographically diverse countries spanning an epidemiologic transition. Overall, our results provide evidence that the gut and oral microbiota may potentially be both predictive as well as a therapeutic target for elevated CM risk, and is in line with previous evidence that the human microbiota are associated with CM risk(40).

Consistent with previous studies(9, 41), we found that the gut microbial alpha diversity was significantly lower in participants with elevated CM risk. However, these associations are mostly geographical and dependent on the type of CM risk factor, e.g. for elevated blood pressure, the association was only found among South Africans and Ghanaians, while only Jamaicans had differences for elevated fasted blood glucose. Similarly, gut bacterial beta diversity was also significantly different between participants with an elevated waist circumference, elevated blood pressure, hypertriglyceridemia or low HDL concentration, compared with their healthy counterparts. Although again, the associations differed by country, e.g. for an elevated waist circumference, the associations were only found among the South African and Ghanaian participants, and for low HDL concentration, only among the US participants.

Previous studies have reported conflicting results as to which specific bacterial taxa associate with CM risk factors(11–13, 17, 18). In our study, gut bacterial ESVs annotated to family Lachnospiraceae were significantly enriched in participants with the elevated CM risk phenotype, along with individual risk factors, including elevated waist circumference and hypertriglyceridemia. Previously, a number of studies have found that an enrichment of Lachnospiraceae was associated with the development of obesity, insulin resistance and other metabolic disorders(42–46). Notably, participants with an elevated waist circumference also showed an enrichment for the genus *Streptococcus*, which has previously been found to be enriched in some CM diseases(17). In our study, participants with healthy waist circumferences, and blood pressure and a normal fasting blood glucose exhibited a significant enrichment of the Ruminococcaceae family (supplementary results), which includes several kinds of beneficial bacteria known to be negatively correlated to metabolic disease and associated with lower CM risk, e.g. genus *Faecalibacterium*(47, 48) and genus *Oscillospira*(49). Similarly, gut bacterial ESVs annotated to the Clostridiaceae and Peptostreptococcaceae families and *Prevotella* genus were enriched in participants with a lower total CM risk as well as individual CM risk factors, e.g. healthy waist circumferences and normal fasting blood glucose levels, consistent with previous research, reporting a negative association with obesity or other CM risk factors(18, 50, 51). Although it should also be noted that there are a handful of studies that suggest that Clostridiaceae and *Prevotella* are significantly enriched in patients with metabolic diseases(51–53).

Our predicted metagenomic functional analysis showed that participants with elevated CM risk harbored a more pronounced inflammatory phenotype. Indeed, predicted genes involved in LPS biosynthesis pathways were significantly enriched in participants presenting with individual CM risk factors across diverse geographic settings. Similar findings have been reported previously and suggest increased LPS synthesis potential in the gut microbiota in people with obesity, diabetes and other related metabolic diseases(19, 20). Of note, the predicted relative proportion of genes that encode for enzymes involved in butyrate synthesis pathway suggested that the lysine pathway, glutarate pathway, and 4-aminobytyrate/succinate pathway were significantly enriched in individuals with elevated CM risk and also individual risk factor. Butyrate can be synthesized via different substrates, driven by enzymes that are produced and secreted by different bacteria(54). While the most common pathway for butyrate synthesis is via pyruvate and acetyl-coenzyme A, other less-dominant pathways include amino-acids (lysine, glutarate and 4-aminobytyrate/succinate) as substrates via the 4-aminobutyrate pathway, which can produce pro-inflammatory byproducts(54) and are related with obesity. Together, these results suggest that the gut microbial metabolic functional potential of participants with elevated CM risk had a marked inflammation-driving capacity, which may influence host systemic inflammatory levels and may ultimately lead to the CM disease consequence.

While the impact of ethnicity on the core of gut microbiota has been demonstrated by several studies(55, 56), exploring the impact of ethnicity and the associations between gut microbiota and CM risk has not been performed to date. Some studies find that there is a decrease in the gut microbial alpha-diversity that is associated with CM risk from different ethnicities, e.g. Danish obese individuals when compared to non-obese individuals9 and obese compared with lean in a mid-western US female adolescent twin cohort study ^(41)^. Similarly, previous studies find that there is a shared alteration in the gut microbial composition of the individuals with elevated CM risk and adiposity derived from different ethnicities. For example there was an adiposity-related enrichment of Lachnospiraceae in the gut microbiota of participants enrolled in the Twins UK cohort study(46), as well as among British obese individuals(45), and obese, Colombian subjects (57) with increased cardiometabolic risk, and finally among individuals in the midst of Westernization(58) when compared to the lean individuals, respectively. Interestingly, the association between TMAO and cardiovascular disease has been found to be significantly greater in whites compared to blacks^(24)^. In our study of African-origin adults, there was no significant association between predicted genes involved in TMAO biosynthetic pathway and total CM risk or individual CM risk factors, consistent with previous research, finding a negative association between TMAO concentration and cardiac death in black participants^(24)^.

Numerous reports have implicated a close linkage between oral infections, particularly periodontitis, and several systemic diseases, e.g. atherosclerosis, type 2 diabetes mellitus, atherosclerotic cardiovascular disease(30–32). In our study, we found that the oral microbial structure (measured by beta diversity) was associated with indices of elevated CM risk, e.g. high waist circumference, elevated fasted blood glucose, and low HDL concentration. However, as with gut microbiota, these associations varied according to the geographic region. There were also several ESVs significantly associated with total elevated CM risk as well as the individual CM risk factors. For example, participants with total elevated CM risk or individual CM risk factors, e.g. elevated waist circumference, low HDL concentration, elevated blood pressure or hypertriglyceridemia, exhibited enriched oral bacterial ESVs annotated to genus *Streptococcus, Prevotella*, or *Veillonella*, which were previously reported to be associated with obesity and other cardiometabolic disease(59–62). Systemic inflammation is an important component of the role of oral bacteria in the pathogenesis of systemic diseases(30, 34). Some strains in genera *Streptococcus, Prevotella*, or *Veillonella* are opportunistic pathogens and have been previously reported to be associated with inflammatory diseases(63–65), even though, we also found that ESVs in genus *Neisseria* were higher in participants without individual CM risk, e.g. elevated waist circumference, elevated blood pressure or hypertriglyceridemia, inconsistent with previous reports as described above. Importantly, we found a number of instances whereby different ESVs of the same genus were associated with differential CM risk. For example, different ESVs belonging to the genus *Rothia* were significantly enriched in the oral microbiota of participants both with and without elevated CM risk. While it is possible that these are spurious relationships, it is also possible that these two unclassified *Rothia* strains have different genetic makeup in non-genera conserved regions or different transcriptional and/or translational profiles in genus conserved regions associated with CM risk (e.g., distinct immunological and/or metabolic properties) that could result in different host-associated impact. Indeed, this may explain the conflicting results with certain studies(66). The oral microbiota may significantly impact the CM risk through triggering of inflammatory processes. Consequently, an analysis of the oral microbiota can be imagined to be used as a pre-screening test for CM risk, despite differences between the gut and oral bacterial diversity and taxa associated with elevated CM risk. Notably, in this study, we also found a weak but significant association between the oral and gut microbiota, within the structure and also the functional taxa. Thus, it is conceivable that the oral microbiota might be used as a proxy to determine gut-derived associations with CM risk.

In conclusion, our findings extend our insights into the relationship between the human microbiota and elevated CM risk at the structural and functional level, pointing to possible future therapeutic modalities for CM risk targeting the gut and oral microbiota. Our findings identify previously unknown links between gut and oral microbiota alterations, and CM risk, suggesting that gut and oral microbial composition, structure and predicted functional potential may serve as predictive biomarkers for identifying CM risk, as well as therapeutic targets. The features of association between human microbiome and CM risk is diverse in different populations and indicates that different interventions and potential individualized treatment methods targeting the microbiome need to be developed to control the development of CM risk across the world. Indeed, we are continuing our exploration of these relationships in our large international cohort of African-origin adults spanning the epidemiologic transition(35).

## Methods

### Study participants

#### Participant selection

Previously, 2,506 African-origin adults (25-45yrs), were enrolled in METS between January 2010 and December 2011 and followed on a yearly basis. A detailed description of the METS protocol has previously been published(35). For the current study, fecal samples from 655 men and women from Ghana (N=196), South Africa (N=176), Jamaica (N=92) and the US (N=191) were collected in 2014. In addition, 620 of them also supplied saliva samples. Participants were excluded from the original study if they self-reported an infectious disease, including HIV-positive individuals (South Africa), pregnant or lactating women, and unable to participate in normal physical activities(35). METS was approved by the Institutional Review Board of Loyola University Chicago, IL, US; the Committee on Human Research Publication and Ethics of Kwame Nkrumah University of Science and Technology, Kumasi, Ghana; the Research Ethics Committee of the University of Cape Town, South Africa; the Board for Ethics and Clinical Research of the University of Lausanne, Switzerland; and the Ethics Committee of the University of the West Indies, Kingston, Jamaica. All study procedures were explained to participants in their native languages, and participants provided written informed consent after being given the opportunity to ask any questions.

#### Lifestyle and biochemical measurements

All measurements were made at research clinics located in the respective communities. Weight and height were measured. Participants were asked to provide an early morning fecal sample, using a standard collection kit (EasySampler stool collection kit, Alpco, NH) at their home. Fecal samples were immediately brought to the site clinics and stored at −80 °C. Participants were requested to fast from 8 pm in the evening prior to the clinic examination, during which fasting capillary glucose concentrations were determined using finger stick (Accu-check Aviva, Roche).

### Cardiometabolic risk

We defined CM risk using the National Cholesterol Education Program’s Adult Treatment Panel III NCEP/ ATP III criteria for cardiometabolic disease(6). These 5 risk factors include: 1) waist circumference >102 cm in males and >88 cm in females; 2) elevated blood pressure (≥130/85 mm Hg), or receiving treatment; 3) hypertriglyceridemia (≥ 150 mg/dL), or receiving lipid-lowering treatment; 4) low high-density lipoprotein (HDL) cholesterol (<40 mg/dL in males and <50 mg/dL in females), or receiving lipid-lowering treatment; and 5) elevated fasting plasma glucose (≥ 110 mg/dL) or receiving glucose-lowering treatment.

### DNA isolation and 16S ribosomal RNA (rRNA) gene sequencing

Microbial genomic DNA was extracted from the human stool samples using the DNeasy PowerSoil DNA Isolation Kit (Qiagen). The V4 region of 16S rRNA gene was amplified and sequenced using the Illumina MiSeq platform(67). The primers used for amplification (515F-806R) contain adapters for MiSeq sequencing and single-end barcodes allowing pooling and direct sequencing of PCR products(68).

### 16S rRNA gene pyrosequencing data preprocessing and analysis

Raw sequences were pre-processed, quality filtered and analyzed using the next-generation microbiome bioinformatics platform (QIIME2 version 2018.6 pipeline) according to the developer’s suggestion(69, 70). We used the DADA2 algorithm(71), a software package wrapped in QIIME2, to identify exact sequence variants (ESVs). Quality control, filtering low quality regions of the sequences, identification and removal of chimera sequences, merging paired-end reads, which yielded the ESV feature table (ESV table). Alpha and beta-diversity analyses were performed in R using the *phyloseq* package(72). Alpha diversity was calculated by Shannon’s diversity index, observed OTUs, and Chao1 diversity(37). Results were adjusted for BMI, age, sex and country. Principal coordinate analysis (PCoA) was performed based on weighted and unweighted UniFrac distances, a method for computing differences between microbial communities based on phylogenetic information(38). Weighted UniFranc considered both ESVs presence and absence and abundance distances, and unweighted UniFrac only considered ESVs presence. Permutational multivariate analysis of variance (PERMANOVA, R function adonis (vegan, 999 permutations)) was used to analyze statistical differences in beta diversity(73). Benjamini–Hochberg false discovery rate (fdr) correction was used to correct for multiple hypothesis testing(74).

The contribution of CM risk factors and environmental factors (sex, age, BMI, sleep, smoke and alcohol consumption) to the overall weighted and unweighted UniFrac dissimilarities in gut and oral microbiota composition was also assessed using PERMANOVA (R function adonis (vegan), 999 permutations), which decomposes the dissimilarity matrix into ‘variance’ explained by each covariate. The obtained R^2^ gives the proportion of variability observed in the entire dissimilarity matrix that can be independently attributed to the studied variables.

For taxonomic comparisons, relative abundances based on all obtained reads were used. We used the QIIME2 plugin “q2-feature-classifier” and the Naïve Bayes classifier(75) that was trained on the metagenome annotation package Greengenes13.8(76) 99% operational taxonomic units (OTUs) full-length sequences to obtain the taxonomy for each ESV. Significantly differential ESVs were determined using the statistical framework called analysis of composition of microbiomes (ANCOM)(71) for two group comparisons. FDR correction was used to correct for multiple hypothesis testing. Results were adjusted for BMI, age, sex and country.

A two-side Mantel test using Spearman correlation coefficients (999 permutations) was applied to identify the correlation between the beta diversity of the oral and the gut microbiota, with both unweighted and weighted UniFrac distance matrices in software R with the function “mantel.test”(77).

To test for correlations between oral microbiota and the Shannon index or ESVs, which their relative abundances was greater than 1% and are also significantly correlated with CM risk in gut microbiota, random forest regression and generalized linear models (GLM) were performed. Random forest regression was done with 1,000 regression trees based on 10-fold cross-validation and performed with QIIME2 plugin “qiime sample-classifier regress-samples” and the Random Forest regressor in the R programming environment. A randomly drawn 80% of samples were used for model training and the remaining 20% were used for validation.

### Metagenome functional predictions of the microbial pathways

We used Piphillin an algorithm to predict the functional profiles of the microbiome (39). Briefly, this algorithm uses direct nearest-neighbor matching between 16S rRNA gene sequencing datasets and microbial genomic databases to infer the metagenomic content of the samples. Gene prediction was performed on ESVs table using online Piphillin (http://secondgenome.com/Piphillin.), with KEGG database version 2017 and 97% identity cut-off. Predicted gene content and gene copy numbers within each genome were retrieved and classified in terms of the ezymes code by each gene as KEGG orthology (KOs)(78). Results were adjusted for BMI, age, sex and country. Statistical analyses were performed in R. Student’s t-test (normally distributed) or Mann-Whitney U test (not normally distributed) was used for to detect differentially abundant KOs between two groups. FDR correction was used to correct for multiple hypothesis testing.

## Acknowledgments

The authors would like to acknowledge the original METS study participants and site-specific clinic staff members. The current work is funded in part by the National Institutes of Health R01DK080763, R01DK090360 and R01DK111848. BTL is supported by the National Institutes of Health under award number, R01DK104927-01A1 and Department of Veterans Affairs, Veterans Health Administration, Office of Research and Development, VA merit (Grant no. 1I01BX003382-01-A1). BPB is supported by the Arnold O. Beckman Postdoctoral Award.

## Contributions

LRD, BTL and JAG supervised the project. LRD, NF, BTL, and JAG conceived the idea. LRD, LL, JPR, and AL, all collected the human data; NF performed the statistical, and computational analyses. NF, LRD, BTL, JAG, BPB, PB, and WK wrote the manuscript. NG (University of Chicago) performed the microbiome sequencing of the human gut microbiota. All authors reviewed the manuscript.

## Supplementary Materials

Supplementary results

Supplementary Figures 1 to 21

Supplementary Table 1 to5

